# In Silico Characterization of a Hypothetical Protein from *Pseudomonas aeruginosa* LESB58: A Structural and Functional Perspective

**DOI:** 10.1101/2025.04.06.647430

**Authors:** Sazzad Hossain, Anisa Jannat

**Author notes:** Corresponding Author: Sazzad Hossain, Department of Biochemistry & Molecular Biology, Noakhali Science & Technology University, Noakhali-3814, Bangladesh.

## Abstract

**Background:** *Pseudomonas aeruginosa* LESB58 is an omnipresent opportunistic bacteria that causes both acute and persistent infections in immunocompromised people. This bacteria is now on the WHO’s red list, meaning that new medicines are desperately needed for treatment. Although the bacterial genome is known, many hypothetical proteins have unidentified activities. Annotating hypothetical proteins can be critical steps toward discovering new druggable targets and treatments. In this work, a hypothetical protein of the *Pseudomonas aeruginosa* LESB58 strain (accession no. CAW29855.1, 284 amino acids) was chosen for in-depth structural and functional research.

**Methods & Results:** The target hypothetical protein’s subcellular location and other physicochemical characteristics were estimated to indicate that it is cytoplasmic. Using bioinformatics tools, it was determined that the target protein had the conserved domain of the phenazine biosynthesis protein family. Multiple sequence alignment of the target protein’s homologous sequence was produced, which helped to create a phylogenetic tree and identify the target protein’s common ancestor. The extended strand was mostly present in the secondary structure produced by PSI-PRED. Using a template protein (PDB ID: 1uok.1.A), a 3D model of the target protein was predicted using the SWISS-MODEL service based on the homology modeling idea. After energy reduction via the YASARA tool, which increases the structure’s stability the SWISS-MODEL structure was evaluated and validated using multiple tools. The RMSD value of 0.347 Å was produced by superimposing the target with the template protein using UCSF Chimera, indicating a dependable three-dimensional structure. PrankWeb server anticipated and visualized the modeled structure’s active site. The target proteins with non-homologous protein against the human proteome and human antitargets are finally predicted by a pipeline builder, suggest that this protein may be a possible therapeutic target.

**Conclusion:** By helping *P. aeruginosa* evade the host’s immune system and form biofilms that contribute to antibiotic resistance and worsen infections, this protein can boost *P. aeruginosa* pathogenicity. The study’s results, which examine both functional and structural features, might help in the creation of novel antibacterial drug targets and *Pseudomonas aeruginosa* LSB58 medications.

## 1. INTRODUCTION

Hypothetical proteins (HPs), many of which lack experimental proof of production, are encoded in a substantial fraction (30–40%) of bacterial genomes. Our knowledge of these proteins’ biological functions is limited since they are frequently left unannotated in protein databases. These proteins must be annotated utilizing bioinformatics technologies, such as functional prediction, structural analysis, and sequence similarity. Because protein structures are conserved, 3D structure comparison offers more accurate insights than sequence-based approaches.

Computational methods are essential in the analysis of proteins, enabling the prediction of their physical and functional characteristics, the identification of potential drug targets, and the elucidation of protein interaction networks. Functional annotation efforts are improved by methods like machine learning, gene co-expression grouping, and phylogenetic profiling.

The work focusses on a putative protein (CAW29855.1) from the Gram-negative, multidrug-resistant bacterium Pseudomonas aeruginosa LESB58, which is linked to devastating infections in healthcare settings. By forming biofilms and secreting virulence agents including elastase and exotoxins, this strain demonstrates resistance. It is frequently found in natural sources like soil and water as well as in clinical settings.

Clinically, P. aeruginosa causes wound infections, urinary tract infections, and lung infections, particularly in people with weakened immune systems. Culture and molecular methods are used for diagnosis, and antipseudomonal antibiotics are used for treatment; however, resistance makes therapy more difficult, highlighting the need for focused research and innovative drug development methodologies. In order to identify possible therapeutic targets using protein interaction analysis, homology modelling, and virulence profiling, this study uses in silico technologies to structurally and functionally characterize essential hypothetical protein. By identifying important biological pathways and therapeutic entry points, the research seek to develop tailored therapy techniques.

## 2. MATERIALS & METHODS

### 2.1 Study Design

The overall research process is illustrated in figure 1.

**Figure 1:**
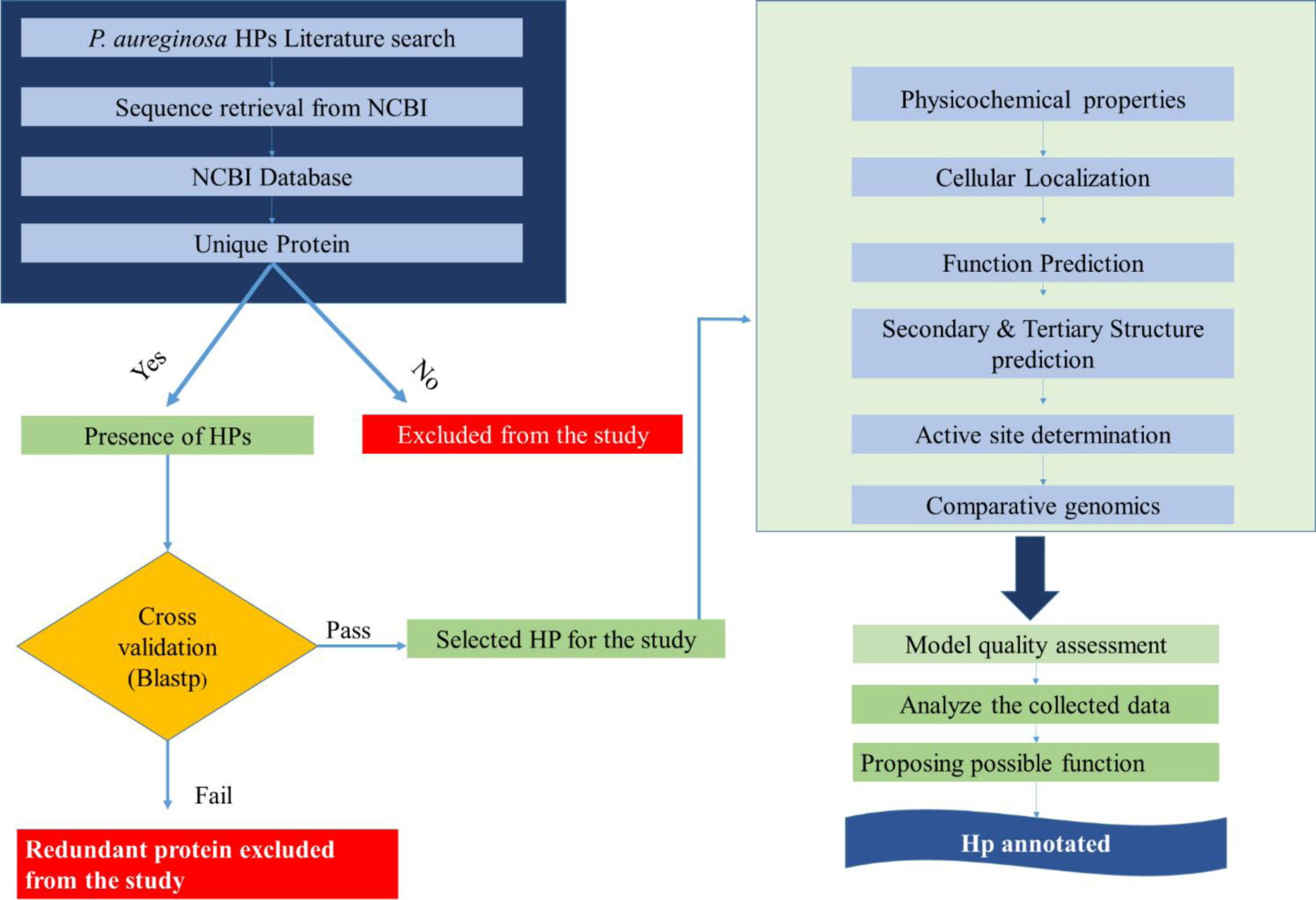
Schematic representation of the whole methodology used in the study.

**Figure 2:**
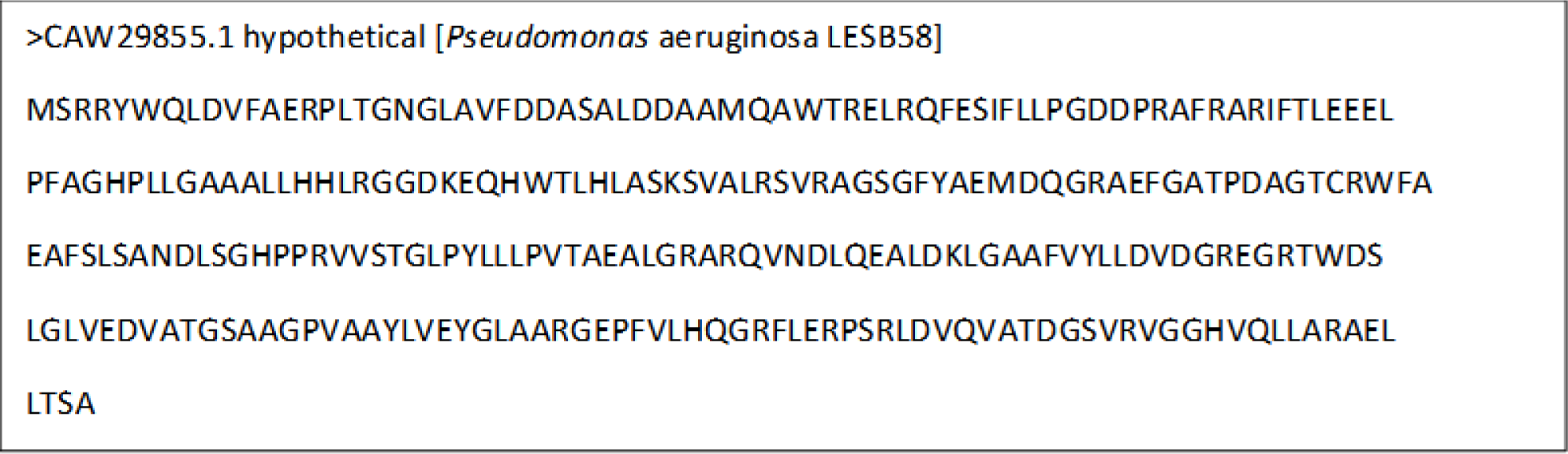
FASTA format from NCBI.

**Figure 2:**
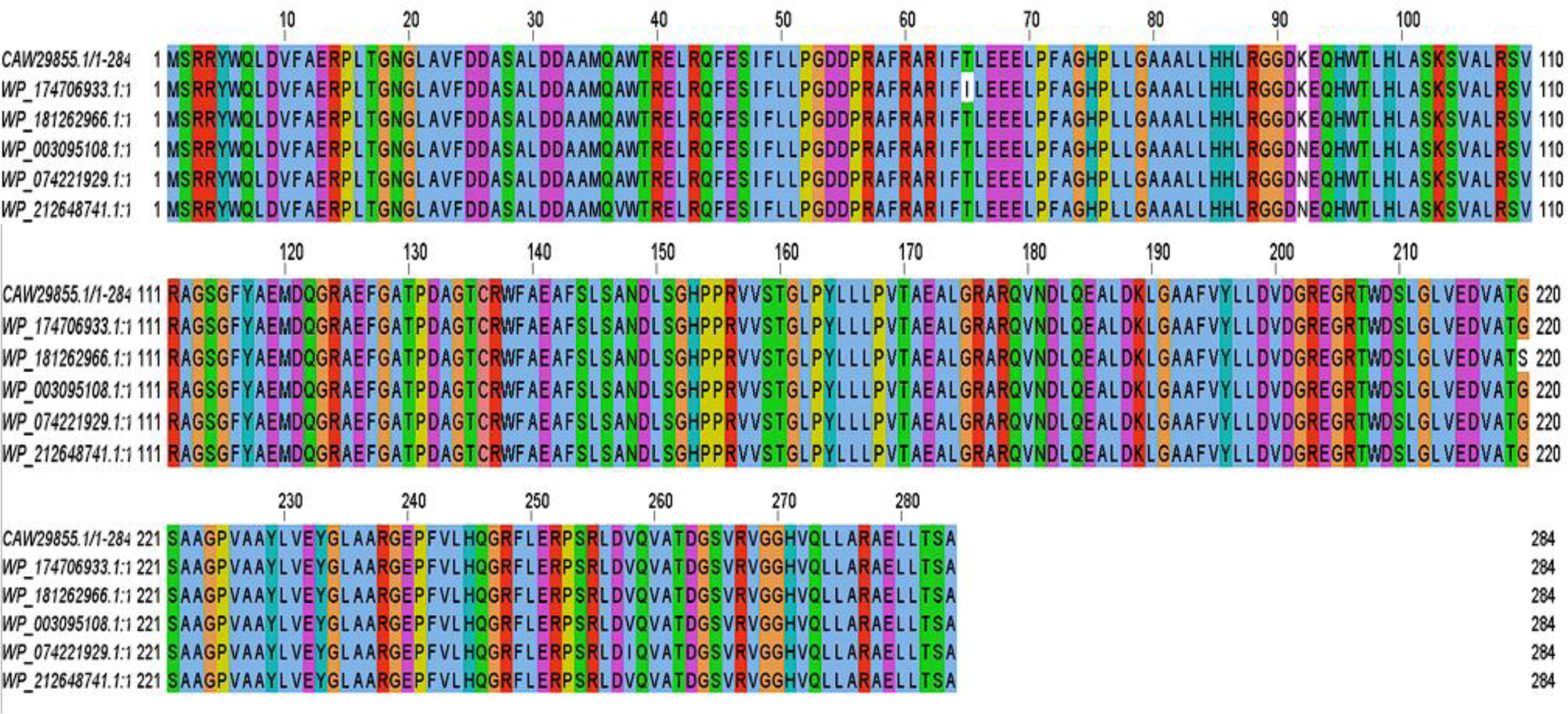
Multiple sequence alignment visualized by Jalview.

### 2.2 Sequence Retrieval

The NCBI database contains 31,087 genomes of Pseudomonas aeruginosa. Genome ASM2664v1, the study used Pseudomonas aeruginosa strain LESB58, which is an NCBI reference sequence. This strain was reported to the Wellcome Trust Sanger Institute in December 2008. This strain has a total chromosomal length of 6.6 Mb. An uncharacterized protein (CAW29855.1) from strain LESB58 with 284 amino acid residues was used for this investigation. The protein’s main sequence was extracted in FASTA (Figure-2) format for further study.

### 2.3 Analysis of Physicochemical Properties

The ProtParam tool from the Expasy web service was utilized to analyze the physicochemical characteristics of a chosen hypothetical protein(23). ProtParam is a tool that calculates several physical and chemical parameters for a user-entered protein sequence. A total of positively and negatively charged residues, molecular weight, theoretical pI, atomic composition, extinction coefficient, estimated half-life, instability index, aliphatic index, and grand average of hydropathicity (GRAVY) are among the parameters(23). The amount of light that a protein absorbs at a certain wavelength is indicated by its extinction coefficient. The instability index estimates a protein’s stability in a test tube. The proportional volume occupied by aliphatic side chains determines a protein’s aliphatic index. The total hydropathy values of all the amino acids divided by the total number of residues in the sequence yield the GRAVY value for a peptide or protein(23).

### 2.4 Subcellular localization and solubility prediction

Assessing the subcellular localization of proteins is essential for comprehending their function and is a critical step in the genome annotation process. The identification and location of adhesion-like intercellular proteins are aided by computer analysis. Subcellular prediction techniques such as CELLOv.2.5, PSORTb v3.0.3 and SOSUI server were used to the hypothetical protein that was chosen from *P. aeruginosa* pathogen.

### 2.5 Function prediction: Protein family, domain and motif analysis

The domain and motif analyses have to be completed before the precise function prediction can be performed. Three bioinformatics tools and databases were used for domain analysis of hypothetical proteins: InterProScan, Pfam, and NCBI Conserved Domain Search Service (CD Search. For motif analysis, use the MOTIF server.

### 2.6 Multiple sequence alignment (MSA) and Phylogenic tree

To find protein homologs, performed a protein BLAST search on the NCBI website using default parameters against the non-redundant database. From the BLAST result 5 identical sequence was downloaded and with the target HP sequence the BLAST sequences were employed to ClustalW to generate multiple sequence alignment. The study used the flexible bioinformatics tool Jalview Version 2 to perform a thorough examination. For Data Alignment and Retrieval Jalview executes multiple sequence alignments (MSAs) after extracting protein sequences from pertinent databases. The study was able to see and modify the alignments with the system, which guaranteed that gaps and preserved areas were accurately represented. Phylogenetic tree was constructed by utilizing the aligned sequence using Jalview. Functional Annotations predicted functional motifs and domains inside hypothetical proteins by utilizing Jalview’s annotation capabilities.

### 2.7 Secondary Structure Determination

The study employed the self-optimized prediction technique with alignment (SOPMA), which may be found at (https://npsaprabi.ibcp.fr/cgibin/npsa_automat.pl?page=/NPSA/npsa_sopma.html) to forecast secondary structure. To verify and validate the results obtained by SOPMA, the study has also used supplemental tools like PSIPRED (http://bioinf.cs.ucl.ac.uk/psipred/).

### 2.8 3D structure modeling

Template-based homology modelling was used to establish the 3D structure of the protein. The SWISS-MODEL service, which creates homology models by aligning the target sequence with the template structure using template search, was used to produce the three-dimensional structure of the targeted protein. In order to build the model as accurately as low-resolution X-ray crystallography, I only take into account templates that have at least 30% sequence identity. I-TASSER web tool was also used to determine the 3D model of the target HP. Lastly, Galaxy Refiner was used to optimize the constructed structures; the model that produced the best results was determined by finding the lowest MolProbity and highest GDT-HA value. Consequently, the revised structure files, which are in the.pdb format, were visualized using BIOVIA Discovery Studio Visualizer (version 20.1.0.19295).

### 2.9 Energy minimization of the model structure

The 3D model architecture produced by the SWISS-MODEL & I-TASSER server was subjected to YASARA force field reduction for energy optimization. The target protein is represented in three dimensions more accurately and steadily after this process, which also increases structural accuracy.

### 2.10 Quality assessment

The quality of the model structure was evaluated using tools such as PROCHECK, Verify3D QMEAN from SWISS-MODEL Workspace, and ERRAT. Likewise, UCSF Chimaera software was used to visualize the model and template structures. On top of that, the Z scores for both proteins were calculated using the ProSA-web server.

### 2.11 Molecular Dynamics Simulation

A user-friendly, web-based interface was employed in the study to propose the predicted protein architectures. Here, I predicted the structure of the desired protein candidate using template-based homology modeling. I simulated protein in the water dynamics simulation for more than 50 ns using the WebGro GROMACS simulation tool. Preprocessing, energy reduction, equilibration, molecular dynamics, trajectory analysis, and result production are some of the processes that the simulation system goes through to operate. The radius of gyration (Rg), solvent-accessible surface area (SASA), hydrogen bonds, root mean square deviations (RMSD), and root mean square fluctuation (RMSF) were all measured using the trajectory analysis.

### 2.12 Active site determination

PrankWeb service was utilized to determine the protein’s active site. PrankWeb is a web-based tool for predicting and visualizing protein-ligand binding sites. It relies on P2Rank, a machine learning-based technique for predicting ligand binding sites from protein structures1. The tool’s main features include predicting and visualizing protein-ligand binding sites, as well as comparing the location of predicted pockets with highly conserved areas and actual ligand binding sites. Because of this, PrankWeb is now a crucial tool for identifying the precise locations and important residues of proteins that engage in ligand interactions. PyMOL, an acknowledged bioinformatics application, is used to investigate and describe putative proteins (32). This in silico method used many important PyMOL applications: First, the study used PyMOL, a molecular viewer, to see tiny macromolecules, including proteins. The study has been able to investigate minute aspects of protein structures, such as solvent accessibility, ligand binding sites, and secondary structures. Secondly, PyMOL made it possible to work with and examine protein structures. Particular residues, domains, and interactions were emphasized in the study. Structure-related research was facilitated by visualization techniques including surface representations and ribbon diagrams (32). The PrankWeb findings were also visualized using PyMOL software.

### 2.13 Comparative genomics approach: Non-homology identification

To explore whether the hypothetical protein CAW29855.1 shares any similarities with humans, the current research conducted a BLASTp search against the Homo sapiens proteome. Any human disease’s medication target has to be identified using a few characteristics. Because of this, the study attempted to analyze non-homology characteristics by using the pipeline builder for the identification of target (PBIT) server for non-homology analysis against the human anti-targets and proteome. The work began by identifying human homologous proteins that have a high degree of sequence similarity with the human proteome using the pipeline builder. With the E − value > 0:005 and %sequence identity < 50 set, the BLAST method was used to determine the sequence similarity of the inputted HP sequence. Following that, the target HP was applied in the pipeline to identify non-homologous proteins against human anti-targets—proteins that have negative effects due to a medication known as an anti-target. The PBIT database uses the BLAST method, which employs human anti-target proteins based on various literature, to filter out the significant comparable sequences with known human anti-targets. %sequence identity < 50 and E − value > 0:005 were again set.

## 3. RESULTS

### 3.1 Physicochemical properties

The study looked for the physicochemical parameters of research HP. The proteins had a molecular weight of 30837.79 Dalton. The theoretical pI (isoelectric point) refers to the pH at which the charge of an amino acid in a protein stays neutral. As a result, no movement occurs when put in an electric field with a direct current. Proteins are dense and stable at isoelectric pH, which makes this parameter useful. The theoretical pI range is computed as 5.28. Both metrics (molecular weight and theoretical pI) aid in the visualization of two-dimensional gel electrophoresis, or (2-DE), and hence contribute to scientific studies of these hypothetical proteins. The aliphatic index is a useful indication for measuring the thermo-stability of particular protein molecules. The protein’s tabulated aliphatic index was 92.18. A protein’s stability in a test tube is determined by a metric known as the instability index. We used a threshold value of 40 for our investigation, meaning that a protein is considered stable if its value is less than 40 and unstable if it is greater than 40. The investigation HP shown stability, with an instability index score of 30.02. The amount of protein-water interaction is measured by calculating the grand average of hydropathy (GRAVY), which is computed by dividing the sum of the hydropathy values of all the amino acids by the total number of residues in the given sequence. A protein interacts with water more when its GRAVY value is lower. The research protein’s GRAVY value of -0.074 indicated that it was interactive with water. Physicochemical properties are shown in Table 2.

**Table 1:** Bioinformatics resources used in the study.

| Server/database | Using reason | Link |
| --- | --- | --- |
| ProtParam | Computation of various physical and chemical parameters for a given protein | https://web.expasy.org/protparam/ |
| CELLO | Subcellular localization | http://cello.life.nctu.edu.tw/ |
| TMHMM | Prediction of transmembrane helices in proteins | http://www.cbs.dtu.dk/services/TMHMM-2.0/ |
| PROTTER | Predicts the presence and location of signal peptide cleavage sites in amino acid sequences and prediction of transmembrane helices in proteins | https://wlab.ethz.ch/protter/start/ |
| PSIPRED | Secondary structure prediction | http://bioinf.cs.ucl.ac.uk/psipred/ |
| I-TASSER | Secondary structure prediction | https://zhanggroup.org/I-TASSER/ |
| SWISS-MODEL | Protein 3D structure determination | https://swissmodel.expasy.org/ |
| Galaxy refine | Refinement of protein structure | http://galaxy.seoklab.org/cgi-bin/submit.cgi?type=REFINE |
| PrankWeb | Active site determination | https://prankweb.cz/ |
| PyMOL software | Structure visualization | https://pymol.org/2/ |
| ERRAT | 3D structure validation | https://saves.mbi.ucla.edu/ |
| Verify3D | 3D structure validation | https://saves.mbi.ucla.edu/ |
| PROCHECK | 3D structure validation | https://saves.mbi.ucla.edu/ |
| Ramachandran plot | 3D structure validation | http://services.mbi.ucla.edu/SAVES/Ramachandran/ |
| PBIT | Pipeline building for non-homology analysis | http://www.pbit.bicnirrh.res.in/ |

**Table 2:** Physicochemical properties of the hypothetical protein.

| Parameters | Value |
| --- | --- |
| MW (Dalton) | 30837.79 |
| Theoretical pI | 5.28 |
| Formula | C1378H2141N393O406S4 |
| Total number of positively charged residues (Arg + Lys) | 27 |
| Total number of negatively charged residues (Asp + Glu) | 37 |
| Estimated half life | 30 hours (mammalian reticulocytes, in vitro).<br>>20 hours (yeast, in vivo).<br>>10 hours (Escherichia coli, in vivo). |
| Instability index (II) | 30.02 |
| Aliphatic index | 92.18 |
| Grand average of hydropathicity (GRAVY) | -0.074 |

### 3.2 Subcellular localization & solubility analysis

The study HP’s cellular localization was determined using the CELLO & PSORTb website. According to CELLO data, proteins are located in the cytoplasm (Figure 3). PSORTb server also indicated the protein cytoplasmic. The existence of the transmembrane helix was also discovered, which can aid in the function of a protein via transmembrane transport. The existence of signal peptides was also explored using the websites Signal4.1 and PROTTER. The study HP does not contain any signal peptides. SOSUI server predicted the protein to be a soluble protein.

**Figure 3:**
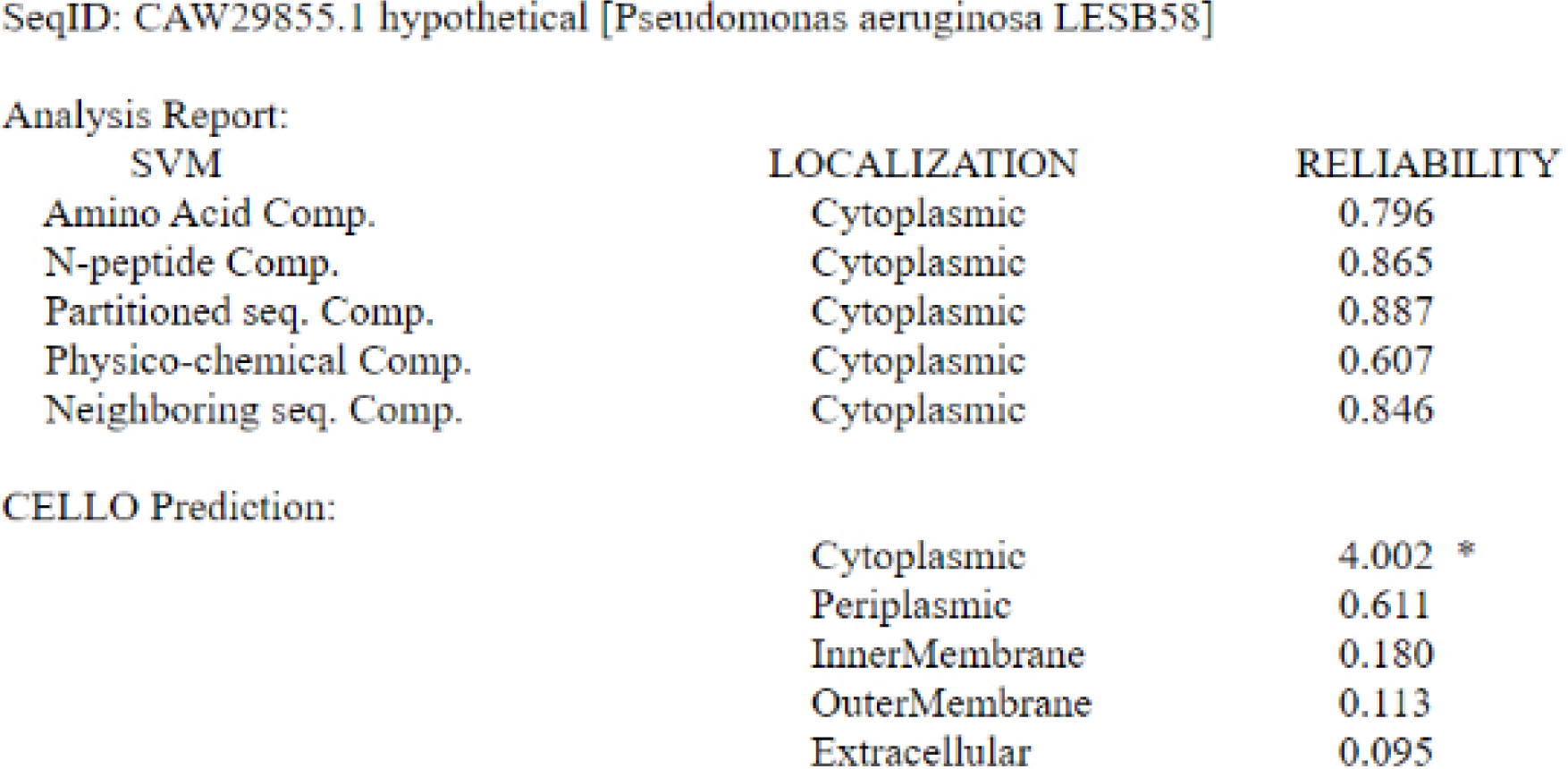
CELLO result of subcellular localization.

**Figure 3:**
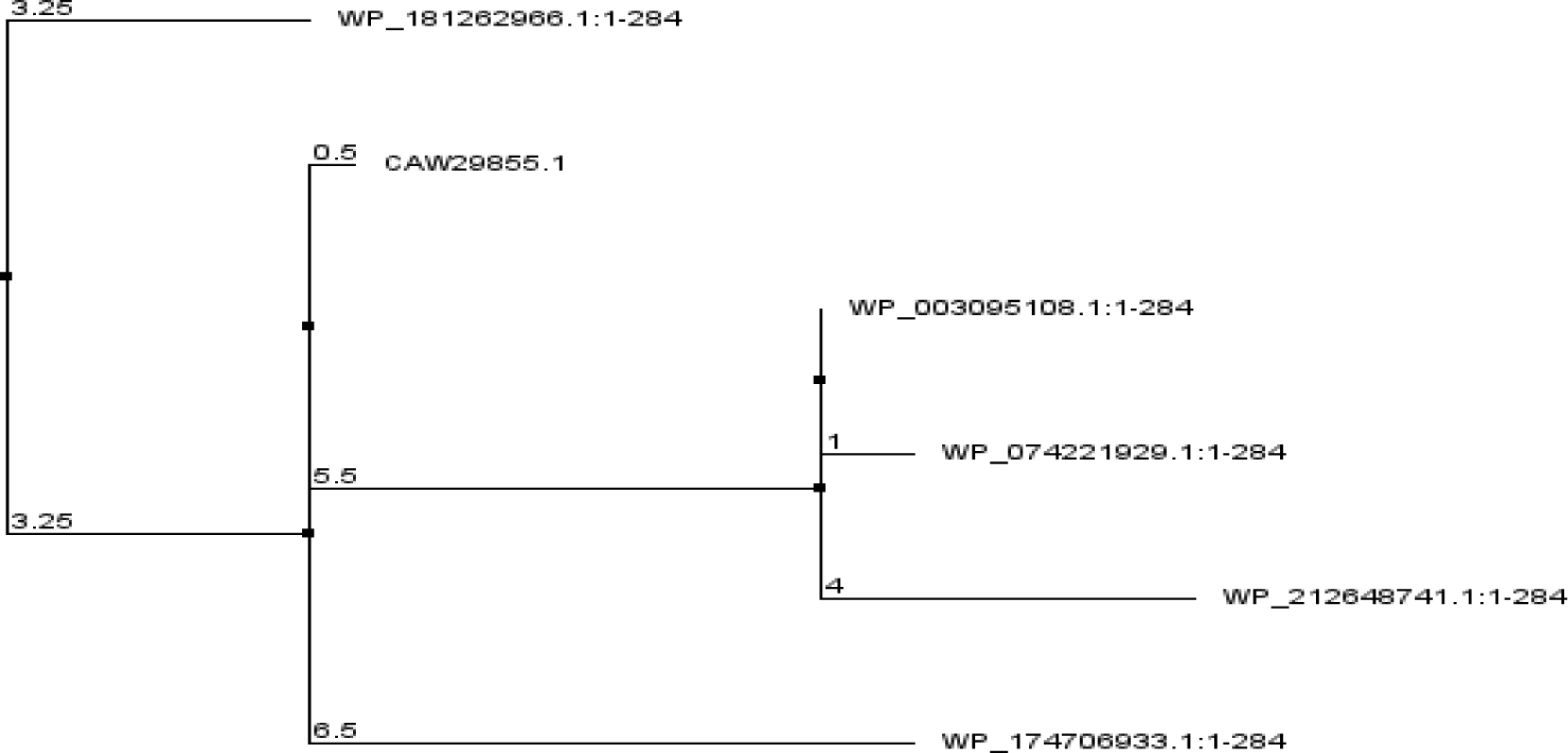
Phylogenetic tree.

### 3.3 Protein family analysis

The study used different annotation techniques to discover conserved domains and putative activities of the target protein. According to predictions made by NCBI-CD Search and InterProScan, the target protein has a domain from the YH19 superfamily, protein family membership of Phenazine biosynthesis PhzF-like. Also predicted catalytic activity as a molecular function and was involved in biosynthetic processes by InterPro GO terms. The NCBI-CDD service found the superfamily domain, which spans amino acid residues 1 to 281, with an E-value of 8.78e-69(Table 6). InterPro’s prediction also placed the superfamily domain between amino acid residues 3 and 282(Table 3). MOTIF analysis verified the existence of a Phenazine biosynthesis-like protein domain from positions 8 to 271, with an E-value of 8.2e-33(Table 3).

**Table 3:** Functional annotation results of different tools.

| Tool name | Predicted domain | Interval | E-value |
| --- | --- | --- | --- |
| NCBI-CDD | YH19 superfamily(Phenazine biosynthesis-like protein) | 1-281 | 8.78e-69 |
| InterProScan | YH19 superfamily(Phenazine biosynthesis-like protein) | 1-282 | 8.67e-69 |
| MOTIF | YH19 superfamily(Phenazine biosynthesis-like protein) | 8-271 | 8.2e-33 |

### 3.5 Multiple sequence alignment and phylogeny analysis

The non-redundant database’s BLASTp search revealed homology with highest sequence similarity to the known PhzF protein from bacterial sources. Five of these sequences that were acquired via BLASTp were then subjected to a multiple sequence alignment using ClustalW & Jalview 2.11.1.3. (Figure 4) presents the alignment findings. Using the same sequences, a phylogenetic tree was created using Jalview software to show their evolutionary relationships, as shown in (Figure 5). The evolutionary links between several phage tail genes are clarified by this graphical depiction, which faithfully depicts the genetic distances.

**Figure 4:**
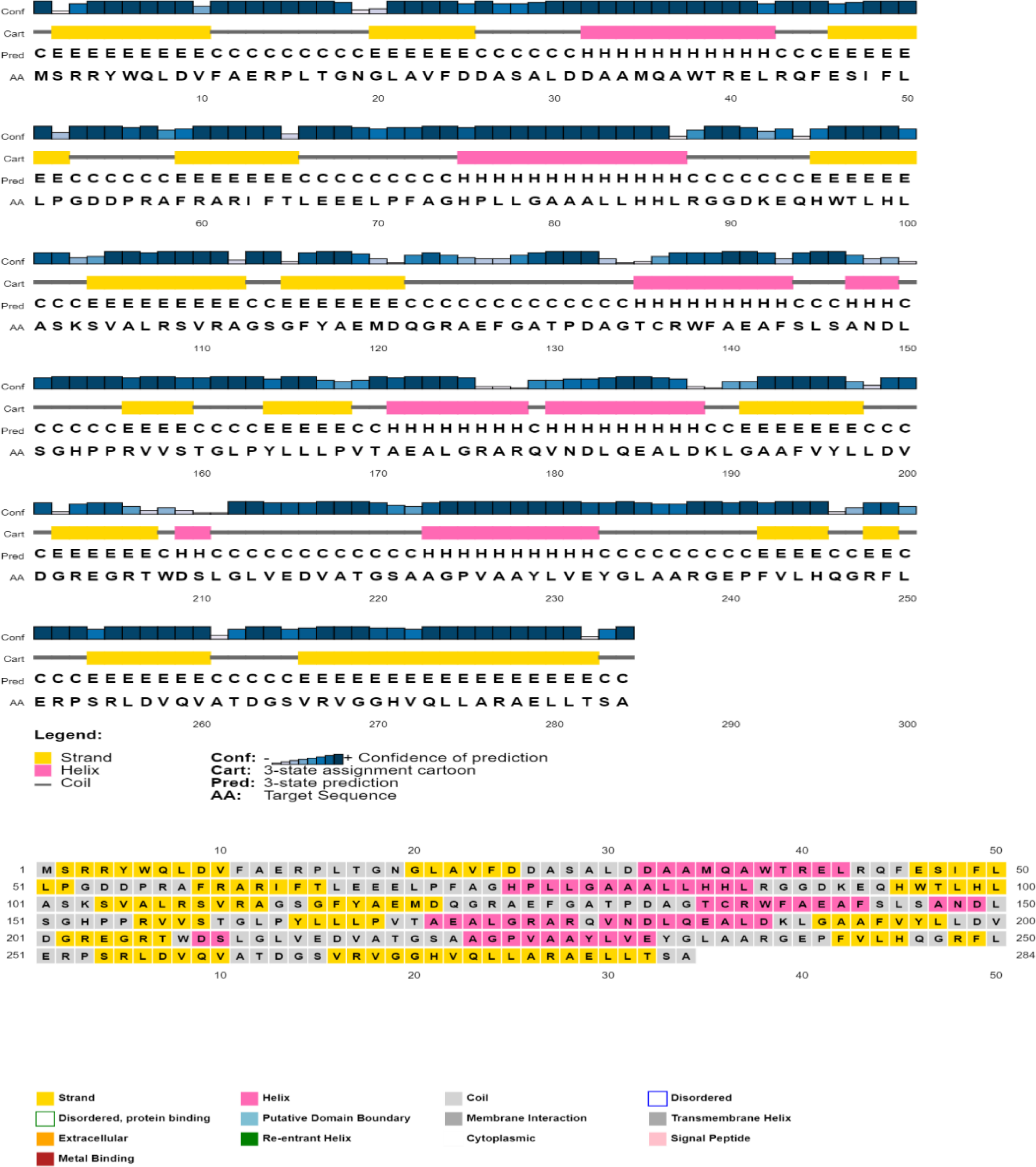
Predicted secondary structure of the target protein using PSI-PRED server.

**Figure 5:**
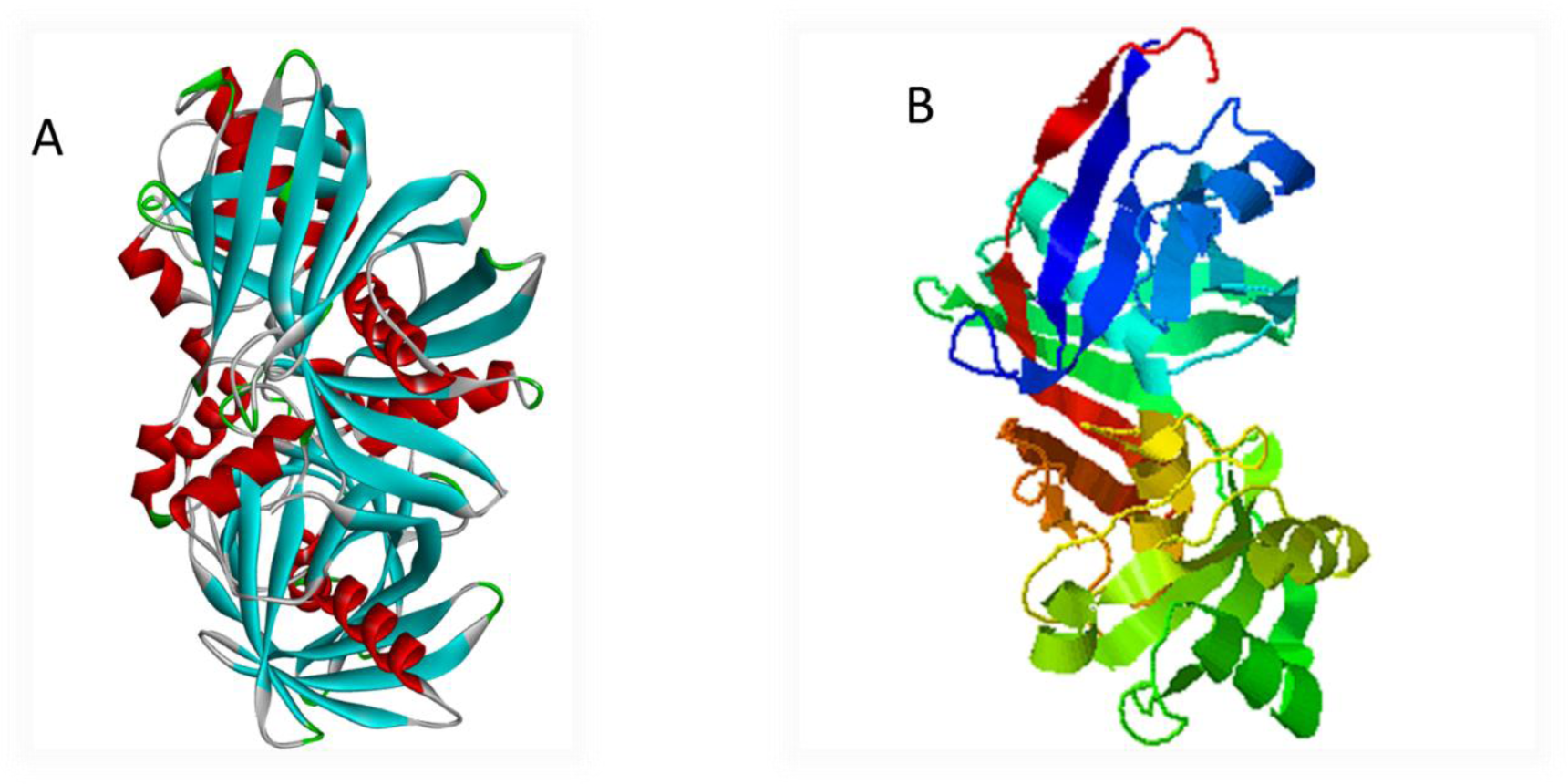
Predicted 3-dimensional structure of the target protein through SWISS-MODEL server after YASARA energy minimization (A), predicted 3D model of the target protein through I-TASSER server (B)

### 3.6 Prediction & analyzing secondary structure

The PSIPRED sequence plot and PSIPRED cartoon plot were provided as a result of the PSIPRED web servers. The sequence plot and cartoon plot structure described that goldenrod (semi-yellow) color is for the extracellular strand domain, pink color is for helix, grey color is for coil, and blackish blue is for the confidence of the structure (Figure 6). According to this, CAW29855.1 showed more strand than helix in its secondary structure. Analyzing the secondary structure through YASARA, the secondary structure of CAW29855.1 content 21.6% helix, 36.5% sheet, 9.8% turn, 32.1% coil, 0.0% 3-10 helix and 0.0% pi-helix.

**Figure 6:**
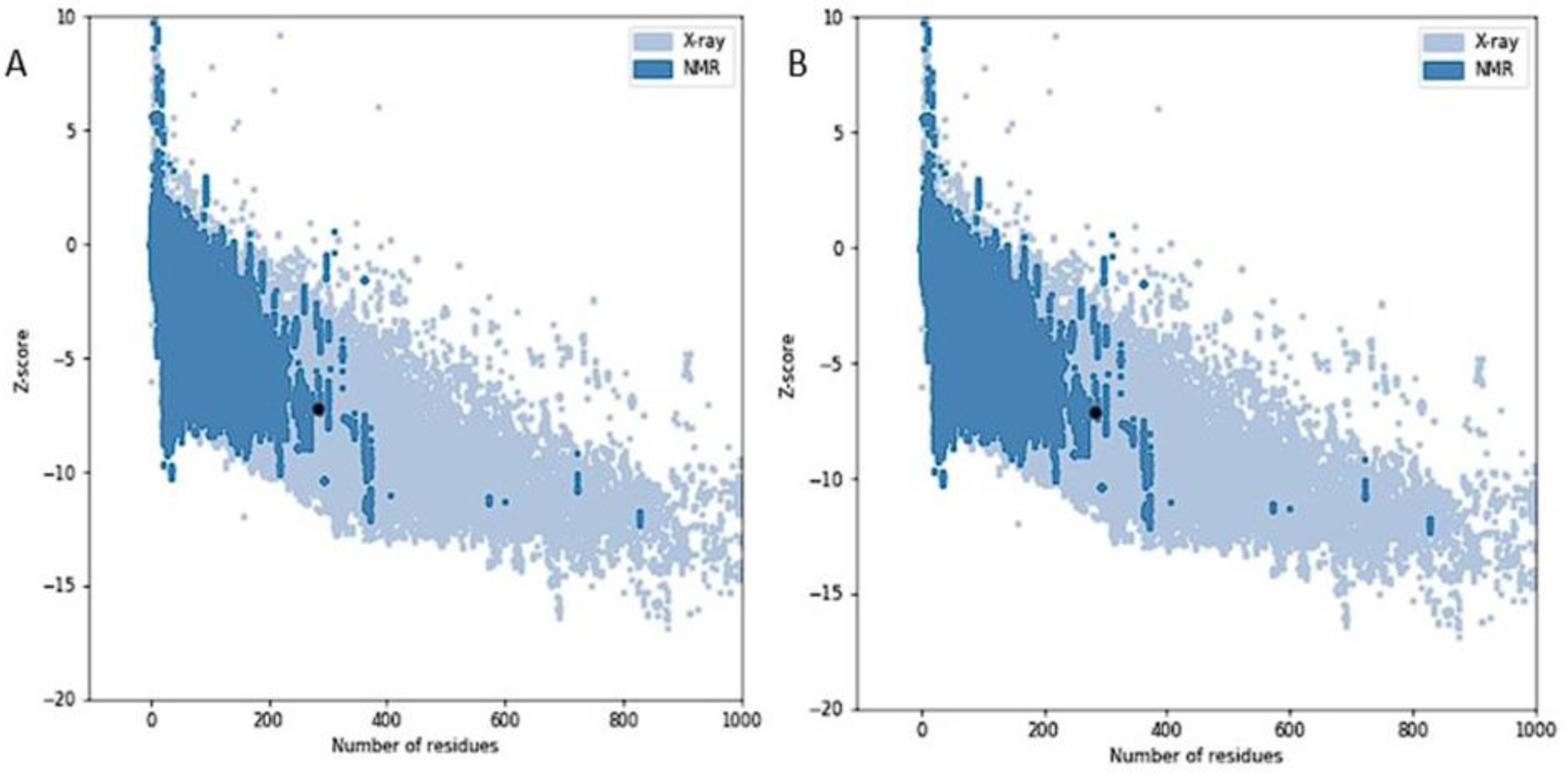
Z scores of the target (A) and template (B) protein using ProSA server. (A: structure from SWISS-MODEL, B: 1uOk.1.A)

### 3.7 3D structure prediction

The 3D structural conformation of the studied protein was determined using SWISS-MODEL’s template-based homology modelling. For template-based homology modeling, look for templates from SWISS-MODEL that match the research protein. 1u0k.1.A gene product PA47161 was chosen for the research HP. The template was chosen based on a number of characteristics, including the Global Model Quality Estimation (GMQE), Qualitative Model Energy Analysis (QMEAN), Z-score, sequence identity, sequence similarity, sequence coverage, and template oligo-state. The template number 1u0k.1.A was a 1.50 Å resolution X-ray diffraction crystallographic structure of *Pseudomonas aeruginosa’s* predicted epimerase PA4716. This template has 99.29% sequence similarity with the 284 AA long CAW29855.1 protein. As a result, the structure from SWISS-MODEL was improved using Galaxy Refiner, and model 6 for the study HP was downloaded following final refinement. The research protein had the lowest MolProbity of 1.597 (model 6), with a starting score 1.051. 3D structure prediction by I-TASSER provided 5 models with properties like C score, Estimated TM-score, Estimated RMSD. Model 1 is selected for having better C score (0.88). The YASARA Energy Minimization Server reduced the energy of the SWISS-MODEL HP structure from -275124.0 kJ/mol to -337496.2 kJ/mol. The preliminary score was -0.73, but after energy minimization, the final score was -0.04, indicating a more stable form and I-TASSER HP structure from −126970.1 kJ/mol to −143007.6 kJ/mol; where the preliminary score was -3.30, but after energy minimization, the final score was -0.94, indicating an improved stability of the form. Structures are shown in figure 7.

**Figure 7:**
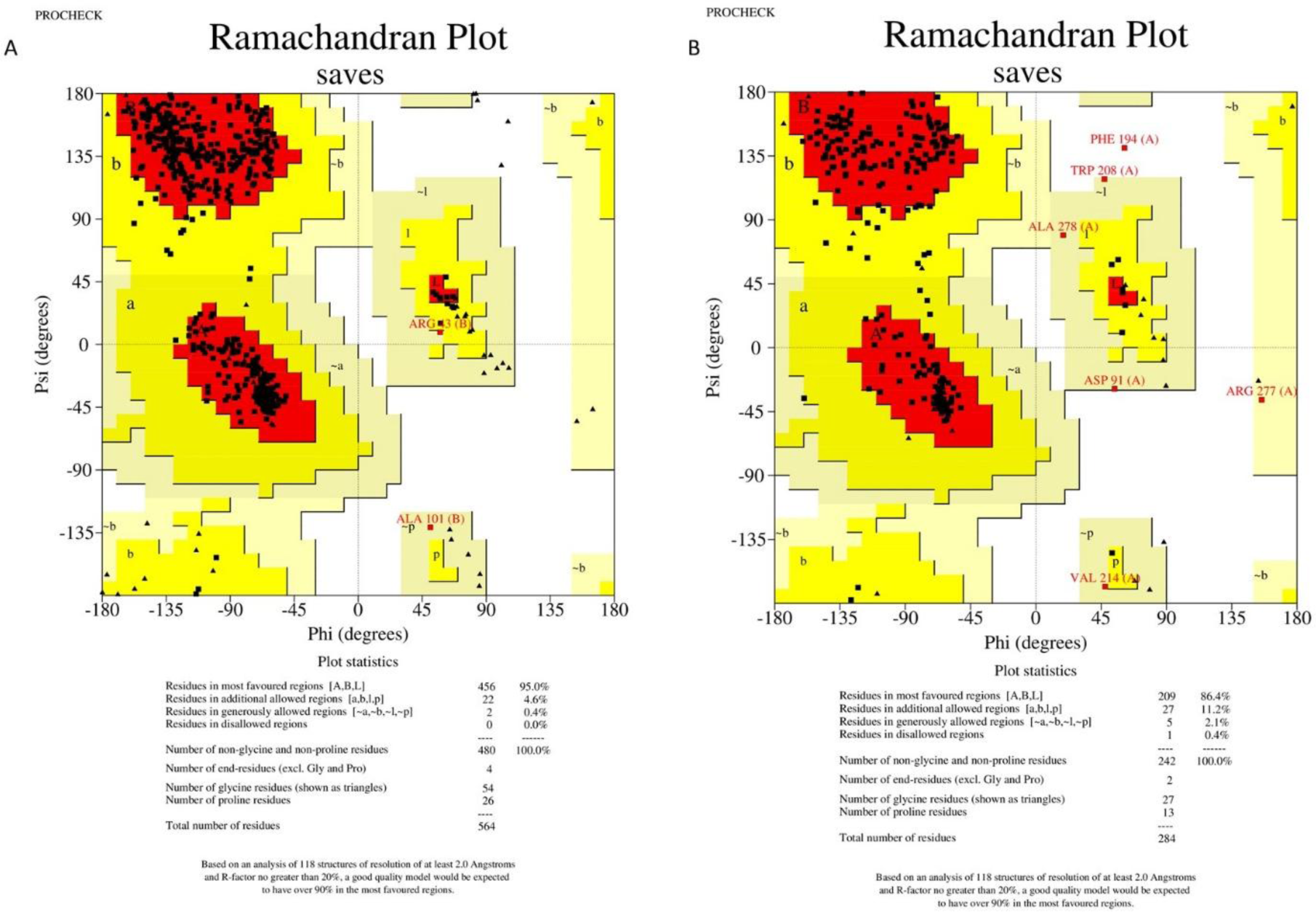
Three-dimensional structure assessment by Ramachandran plot analysis by PROCHECK. (A) Ramachandran plot from SWISS-MODEL (B) Ramachandran plot from I-TASSER.

### 3.8 Quality assessment

The SAVES v6.0 server evaluated the predicted protein structure by running many programs simultaneously to assess the construct model’s quality. The ERRAT value served as the model’s overall quality metric. The overall quality factor for the CAW29855.1 protein structure in SWISS-MODEL and I-TASSER was 96.6912% and 89.4928%, respectively (Table 4). VARIFY3D analyses structures and passes them if at least 80% of the amino acids in the 3D/1D profile with score ≥0.1 The CAW29855.1 structure passes this parameter (based on the SWISS-MODEL and I-TASSER models) (Table 7). WHATCHECK had a color box with a number corresponding to 49 distinct criteria, with green, yellow, and maroon denoting OK, caution, and error, respectively. For structures, the general summary report is acceptable (Table 7). Ramachandran plot analysis from the PROCHECK program also demonstrated that 95% of residues were in the most favored region for structures from SWISS-MODEL & 86.4% from I-TASSER, makes SWISS-MODEL structure a reliable one (Table 4, figure 9). The Z score for the model obtained from ProSA server was −7.2 (Figure 8) and for the template was −7.09 (Figure 8), proposing the homology between the template and the model. Based on UCSF Chimera & YASARA analysis, the target protein (CAW29855.1) and template (1u0k_1) have an RMSD of 0.347 Å over 564 aligned residues with 99.29% sequence identity, indicating a reliable 3D model (Figure 10).

**Table 4:** Three-dimensional structure validation of the predicted hypothetical protein from the SAVES v6.0 server.

| **Protein name** | **ERRAT** | **VARIFY 3D** | **PROCHECK** | **Ramachandran plot**<br>**(% residue in the**<br>**most favored**<br>**region)** |
| --- | --- | --- | --- | --- |
| CAW29855.1(SWISS-MODEL) | 96.6912% | 96.10% of the residues have averaged 3D-1D score≥0.1 (Pass) | Out of 8 evaluations<br>Errors: 0<br>Warning: 6<br>Pass: 2 | 95% |
| CAW29855.1 (I-TASSER) | 90.2174% | 97.18% of the residues have averaged 3D-1D score≥0.1 (Pass) | Out of 8 evaluations<br>Errors: 2<br>Warning: 4<br>Pass: 2 | 86.4% |

**Figure 8:**
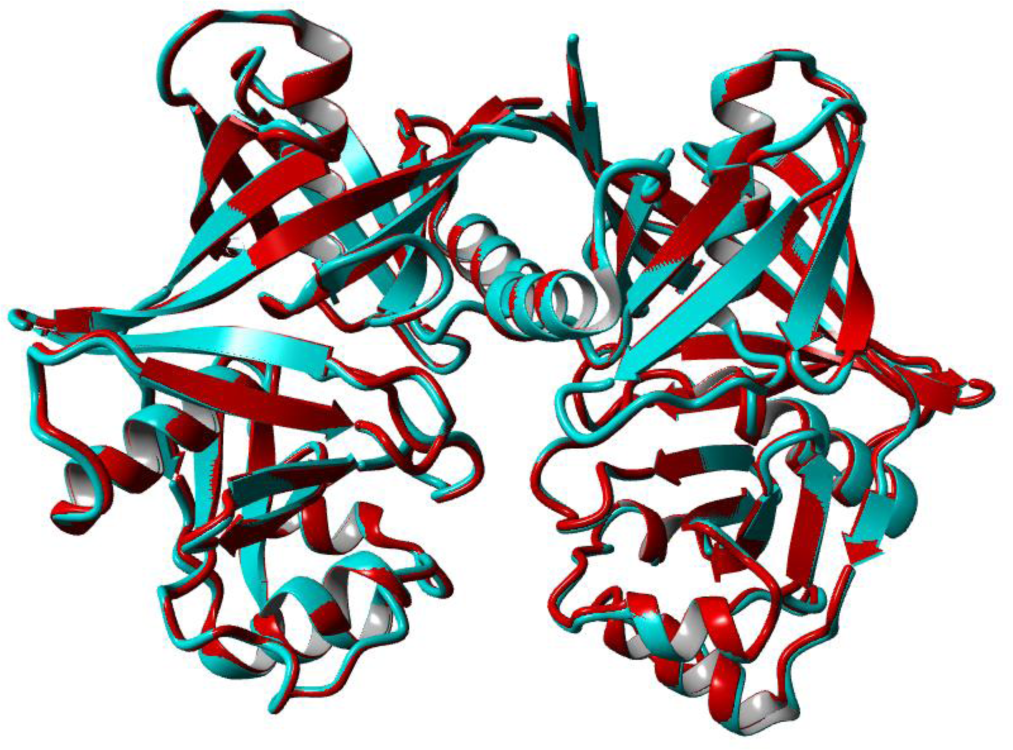
Superimposition of the model (red color) and the template (cyan color) using UCSF Chimera & YASARA tool.

**Figure 9:**
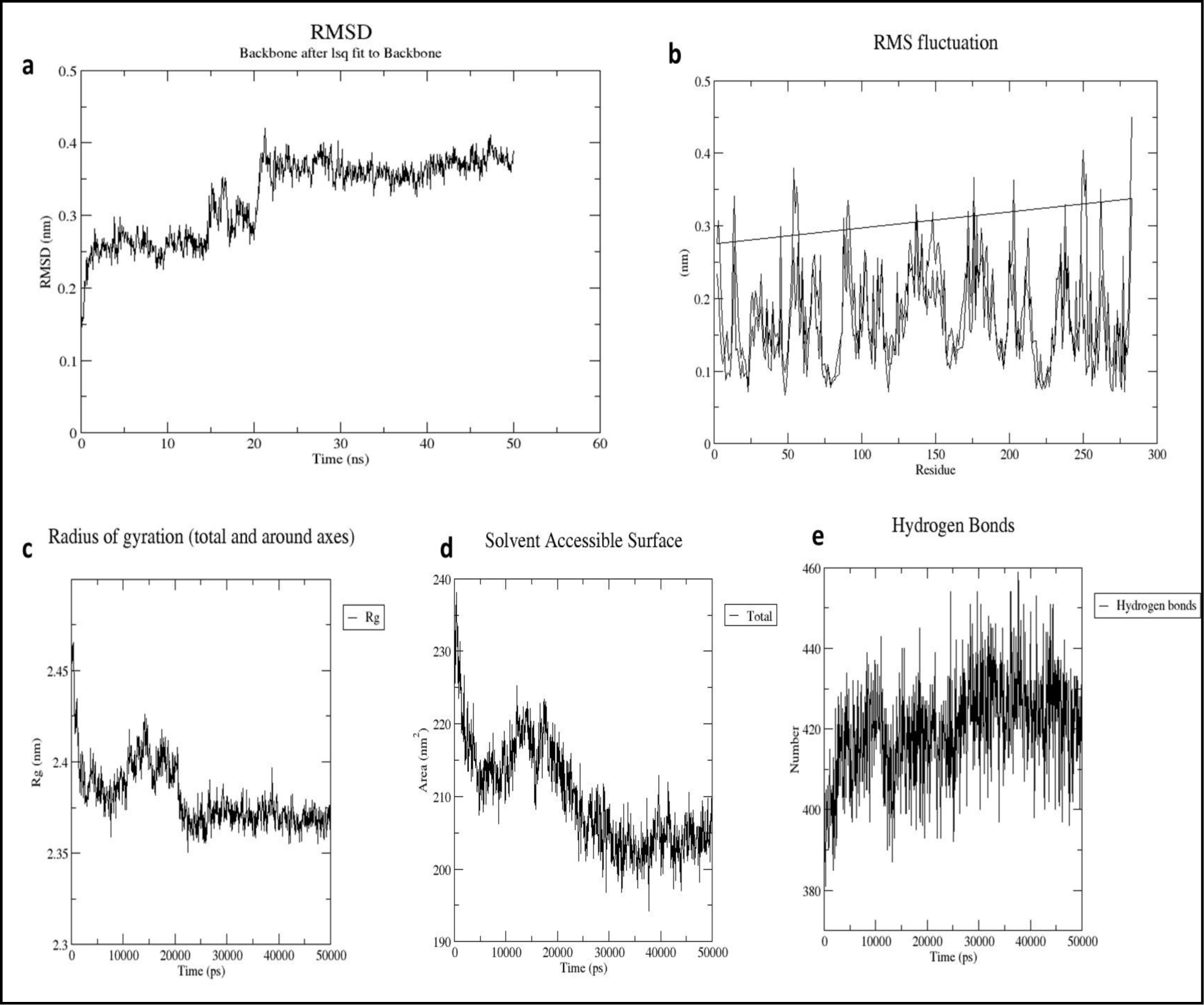
Molecular Dynamics Simulation result (a: RMSD, b: RMSF, c: Radius of gyration, d: Solvent accessible surface area, e: Hydrogen bonds)

### 3.9 Molecular Dynamics Simulation result

As predicted by SWISS-MODEL, the protein structure of CAW29855.1 shows that the RMSD value first rises until 20 ns, stabilizing at 50 ns without experiencing substantial fluctuations (Figure 11a). As a result, the profile of the structure is steady. The RMSF measures the average deviation of amino acid residues during a certain time course. Typically, it gauges the flexibility of each particular residue. Because the structures’ highest RMSF value is less than 0.35 nm, their residue flexibility is smaller (Figure 11b). The analysis of the SASA from the simulation trajectories also shows that the structure predicted by homology modeling has a lower SASA value, indicating a better degree of model stability (Figure 11d). Additionally, it was shown that there is less variation in the proteins predicted by homology modelling by analyzing the Rg value, which is a measure of protein rigidity and mobility (Figure 11c). Over a 50 ns trajectory, the protein structures used in homology modelling exhibit a consistent pattern. In addition, I have ascertained the quantity of hydrogen bonds, which is a crucial factor in determining a stable structure complex. Homology modelling predicts protein structures that have steady hydrogen bonding across a 50 ns trajectory with negligible oscillations (Figure 11e). Throughout the 50 ns simulation trajectory, the overall profile of the protein structure predicted by SWISS-MODEL for CAW29855.1 is stable and constant. Figure 11 displays the outcome of the molecular dynamic simulation.

### 3.10 Active site determination

Based on the simulation results, the SWISS-MODEL structure of the study hypothetical protein was selected for active site prediction. Prank Web assigns pocket scores and probability scores to existing protein pockets. According to the database, the SWISS-MODEL structure of CAW29855.1 included ten pockets. Binding pocket 1 (pocket score: 36.26, probability score: 0.945, AA count: 38); pocket 2 (pocket score: 7.33, probability score: 0.423, AA count: 14); pocket 3 (pocket score: 3.61, probability score: 0.141, AA count: 12); pocket 4 (pocket score: 3.44, probability score: 0.129, AA count: 5); pocket 5 (pocket score: 2.39, probability score: 0.064, AA count: 11); pocket 6 (pocket score: 1.89, probability score: 0.037, AA count: 9); pocket 7 (pocket score: 1.89, probability score: 0.037, AA count: 6); and pocket 8 (pocket score: 1.66, probability score: 0.027, AA count: 5); pocket 9 (pocket score: 1.30, probability score: 0.015, AA count: 6); pocket 10 (pocket score: 0.84, probability score: 0.004, AA count: 6). The protein with binding residue and different color showed different pockets (red for pocket 1; yellow: pocket 2; brown: pocket 3; sky blue: pocket 4; turquoise: pocket 5; blue: pocket 6; pink: pocket 7; green: pocket 8; light green: pocket 9; orange: pocket 10) are shown in Figure 12.

**Figure 12:**
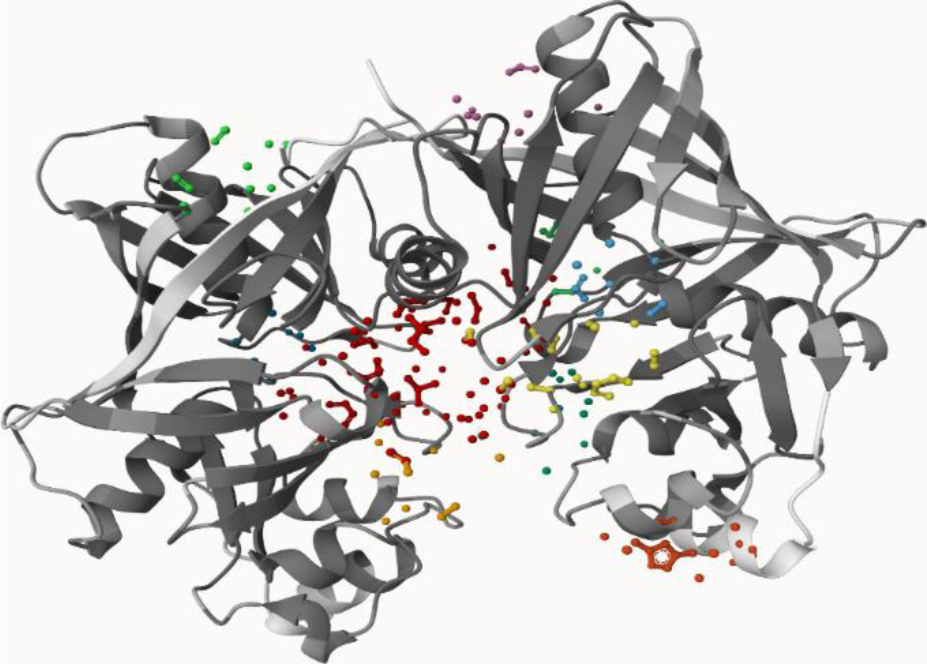
Active site prediction by Prank Web.

### 3.11 Comparative genomics analysis

After completing the structural and functional annotation of the putative protein, the target protein was further characterized using the comparative genomics approach. A new target for a medication must not be identical to human antitargets or the human proteome. The target HP sequence was entered into the pipeline builder from the pipeline builder for identification of the target (PBIT) server to determine the highly comparable sequence with the human proteome. There was no homology between the target HP sequence and the human proteome (Figure 13-A). After that, the sequence was moved to the pipeline analysis that was going to identify the non-homologous proteins that were targeting human antitargets. The target protein is not comparable to the human antitarget, according to this investigation (Figure 13-B). The target protein was identified as a distinct *P. aeruginosa* protein when the data showed no similarity to any known human protein. A good therapeutic option would target microbial proteins that are non-homologous to human proteins to minimize any negative effects.

**Figure 13:**
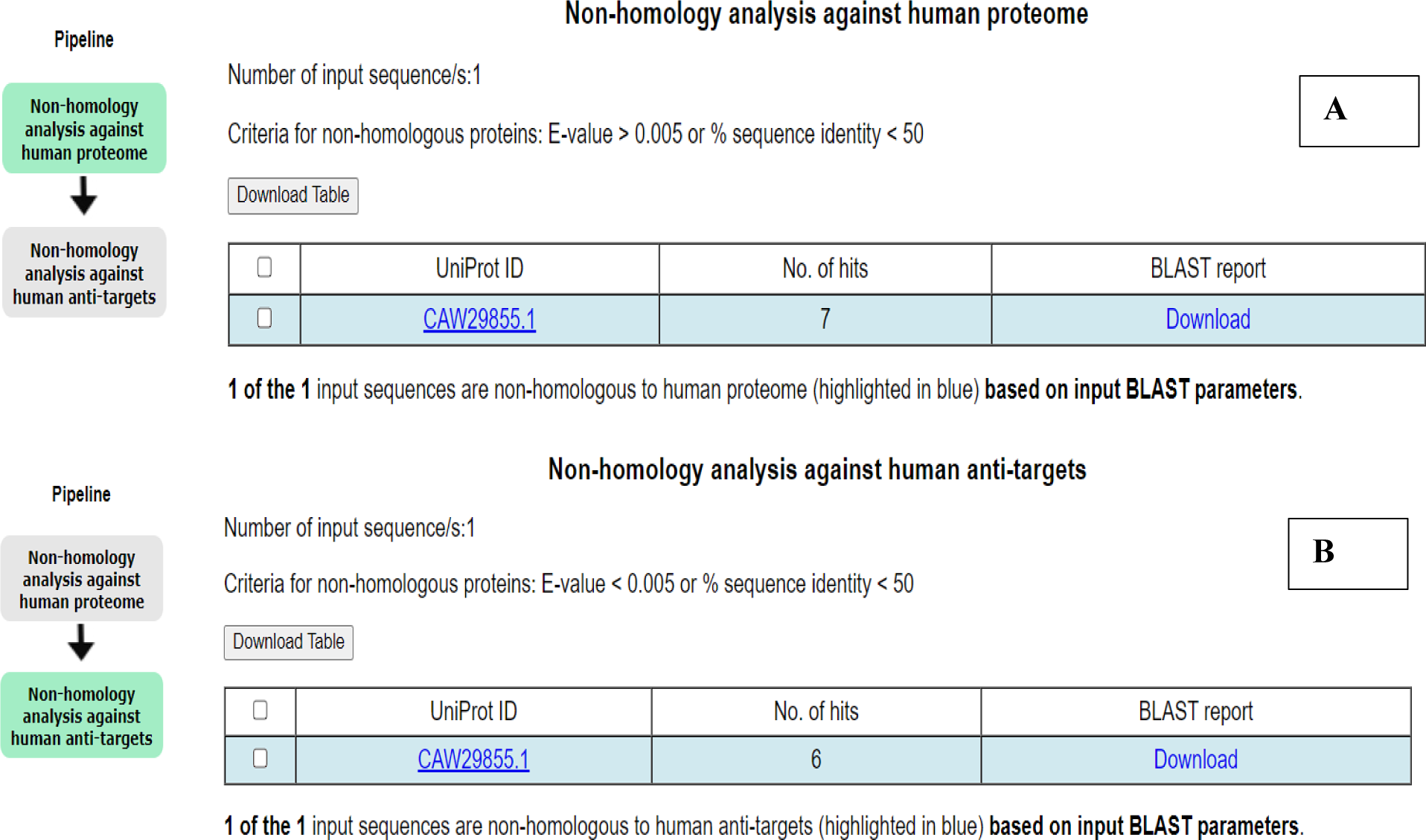
Non-homology analysis against human proteome (A), against human antitargets (B)

## 4. DISCUSSION

Scientists are actively pursuing a vaccine against *Pseudomonas aeruginosa*, but none currently exist. While rapid advancements in affordable gene sequencing have yielded a wealth of genetic and protein data, research on uncharacterized proteins (hypothetical proteins) hasn’t caught up. Understanding these proteins can provide valuable insights into bacterial metabolism, disease progression, drug development, and control strategies. This study employed various bioinformatics tools to analyze the structure and function of a hypothetical protein (CAW29855.1) from the *P. aeruginosa* LESB58 strain. Based on its physicochemical properties, the protein is estimated to be composed of 284 amino acids, with a molecular weight of around 30,8377.9 Daltons, a predicted isoelectric point (pI) of 5.28, and a slightly negative overall hydrophobicity score (GRAVY) of -0.074 (see Table 5). Analysis using CELLO software suggests the protein is located within the cytoplasm (the cell’s interior). The protein’s secondary structure is a mix of random coils, alpha helices, beta turns, and extended strands, with extended strands being the most abundant. All annotation tools confidently predicted the target protein to belong to the YH19/PhzF superfamily, suggesting a role in phenazine biosynthesis. BLASTp result against the nonredundant database showed sequence similarity with other known PhzF family proteins validating the prediction. The generation of phenazines, which are aromatic secondary metabolites containing nitrogen and generated by a variety of bacterial species, depends on the PhzF protein. A crucial step in the production of phenazine is catalyzed by PhzF. It involves two processes in the conversion of (2S, 3S)-2, 3-dihydro-3-hydroxy anthranilate (DHHA): [1, 5]-Hydrogen Shift: This phase exhibits characteristics of both a sigma tropic rearrangement and acid/base catalysis and calls for the conserved glutamate E45. Tautomerization occurs after the [1, 5]-hydrogen shift, producing an aminoketone product1. PhzF coordinates both a water-dependent stage (tautomerization) and a water-free step (the [1, 5]-hydrogen shift). It is a fascinating target for therapeutic research due to its selectivity and crucial function in the production of phenazine.

Three dimensional structure of the protein obtained using SWISSMODEL server successfully passed all of the model quality assessment tools like PROCHECK, Verify 3D, QMEAN and ERRAT. The 3D structure became more stable after YASARA energy minimization process. Superimposition of the model protein with the template protein (PDB ID: 1uOK_1.A) by UCSF chimera & YASARA also suggested the 3D structure to be reliable with RMSD value of 0.347 Å (discussed in “Quality assessment” section). Moreover, from the analysis of molecular dynamic simulation, CAW29855.1 protein structures predicted from the SWISS-MODEL have a consistent and stable profile over the 50 ns simulation trajectory than the other model. The results of the simulation were used to identify the active spots. Furthermore, in order to identify target proteins as potential therapeutic targets, it was necessary to anticipate their active sites in order to determine the likely places at which ligand compounds would bind and start the corresponding processes. The active site amino acid residues computed by PrankWeb server were consistent with the prediction of functional annotation tools and lie in the YH19/PhzF superfamily domain region. Comparative genomics study revealed the protein to be a unique *P. aeruginosa* protein non homologous to human indicating a potential therapeutic target. Further research and experimental validations are needed to confirm our findings about this crucial protein. Phenazines can lead to a variety of diseases in susceptible individuals, including urinary tract infections, burn wounds, pneumonia, and intensive care unit infections. Nosocomial infections can also result from medical equipment infected with biofilm. *P. aeruginosa* presents difficulties due to both its widespread distribution and pathogenic potential. Treatment choices are complicated by antibiotic resistance, and there are few novel medications available. By rupturing bacterial membranes, obstructing electron transport, and producing reactive oxygen species (ROS) inside bacterial cells, phenolazines have antibacterial action. Some phenazines increase the production of biofilms, which increases the persistence of bacteria on surfaces. Bacteria are shielded by biofilms from immunological reactions and environmental stresses. Redox cycling is the process by which phenazines move electrons between their reduced and oxidized forms. ROS are produced by this process, which damages biological components and reduces the survival of bacteria. Phenazines are involved in the pathogenic bacteria’s pathogenicity. The functions of virulence and toxicity factors have been well understood in recent years, but many structural and functional characteristics of other HP and their effectors are still unclear. To the best of our knowledge, this work is the first to describe a *P. aeruginosa* phenazine biosynthetic protein from both a structural and functional standpoint. This hypothetical protein’s annotation might be useful in the development of a successful medication or vaccination. In order to design future treatment options, it will be helpful to comprehend antibacterial processes through the research of individual effectors.

## 5. CONCLUSION

Being acquainted with the functional properties of pathogenic microbes is crucial for biological processes and the study of medicine. The versatile macromolecules known as essential proteins and essential hypothetical proteins can play a critical role in developing novel therapeutic approaches for these dangerous microorganisms. I have combined several bioinformatics databases and tools with an in silico approach to achieve the functional characterization of the target hypothetical protein in this study. I have examined the subcellular location and physiochemical features of the protein and assigned a function to the experiment HP. Moreover, the uniqueness of the protein is predicted by host non homologous analysis. Ultimately, the structural conformation of the HP (CAW29855.1) was ascertained, and an assessment of the predicted model’s correctness revealed that it is a very accurate model. By concentrating on these cutting-edge therapeutic targets, my research will open the door for the development of novel antibacterial medications and therapeutic approaches.

## REFERENCES

1. Uddin R, Jamil FJCb, chemistry. Prioritization of potential drug targets against P. aeruginosa by core proteomic analysis using computational subtractive genomics and Protein-Protein interaction network. 2018;74:115–22.

2. Woods DE, Schaffer MS, Rabin HR, Campbell G, Sokol PAJJocm. Phenotypic comparison of Pseudomonas aeruginosa strains isolated from a variety of clinical sites. 1986;24(2):260–4.

3. Kurowski MA, Bujnicki JMJNar. GeneSilico protein structure prediction meta-server. 2003;31(13):3305–7.

4. Driscoll JA, Brody SL, Kollef MHJD. The epidemiology, pathogenesis and treatment of Pseudomonas aeruginosa infections. 2007;67:351–68.

5. Atron B, Yousif Z. In silico Identification of Novel Therapeutic Targets and Epitopes among the Essential Hypothetical Protein of Pseudomonas aeruginosa: A Novel Approach for Antivirulence Therapy. 2023.

6. Bagag A, Jault J-M, Sidahmed-Adrar N, Réfrégiers M, Giuliani A, Le Naour FJPO. Characterization of hydrophobic peptides in the presence of detergent by photoionization mass spectrometry. 2013;8(11):e79033.

7. Desler C, Suravajhala P, Sanderhoff M, Rasmussen M, Rasmussen LJ. In Silico screening for functional candidates amongst hypothetical proteins. BMC Bioinformatics. 2009;10(1):289.

8. Chakma V, Barman DN, Das SC, Hossain A, Momin MB, Tasneem M, et al. In silico analysis of a novel hypothetical protein (YP_498675. 1) from Staphylococcus aureus unravels the protein of tryptophan synthase beta superfamily (Try-synth-beta_ II). 2023;21(1):135.

9. Wu W, Jin Y, Bai F, Jin S. Pseudomonas aeruginosa. Molecular medical microbiology: Elsevier; 2015. p. 753–67.

10. Poole KJFim. Pseudomonas aeruginosa: resistance to the max. 2011;2:65.

11. Breidenstein EB, de la Fuente-Núñez C, Hancock REJTim. Pseudomonas aeruginosa: all roads lead to resistance. 2011;19(8):419–26.

12. Harris AD, Jackson SS, Robinson G, Pineles L, Leekha S, Thom KA, et al. Pseudomonas aeruginosa colonization in the intensive care unit: prevalence, risk factors, and clinical outcomes. 2016;37(5):544–8.

13. Alhazmi AJIJoB. Pseudomonas aeruginosa-pathogenesis and pathogenic mechanisms. 2015;7(2):44.

14. Streeter K, Katouli M. Pseudomonas aeruginosa: a review of their pathogenesis and prevalence in clinical settings and the environment. 2016.

15. Strateva T, Yordanov DJJomm. Pseudomonas aeruginosa–a phenomenon of bacterial resistance. 2009;58(9):1133–48.

16. Bodey GP, Bolivar R, Fainstein V, Jadeja LJRoid. Infections caused by Pseudomonas aeruginosa. 1983;5(2):279–313.

17. Bassetti M, Vena A, Croxatto A, Righi E, Guery BJDic. How to manage Pseudomonas aeruginosa infections. 2018;7.

18. Rahman A, Sarker MT, Islam MA, Hossain MU, Hasan M, Susmi TFJBRI. Targeting essential hypothetical proteins of Pseudomonas aeruginosa PAO1 for mining of novel therapeutics: an in silico approach. 2023;2023(1):1787485.

19. Robert X, Gouet PJNar. Deciphering key features in protein structures with the new ENDscript server. 2014;42(W1):W320–W4.

20. Reynolds D, Kollef MJD. The epidemiology and pathogenesis and treatment of Pseudomonas aeruginosa infections: an update. 2021;81(18):2117–31.

21. Rahman MA, Heme UH, Parvez MAKJPo. In silico functional annotation of hypothetical proteins from the Bacillus paralicheniformis strain Bac84 reveals proteins with biotechnological potentials and adaptational functions to extreme environments. 2022;17(10):e0276085.

22. Reem A, Zhong Z-H, Al-Shehari WA, Al-Shaebi F, Amran GA, Moeed YA, et al. Functional annotation of hypothetical proteins related to antibiotic resistance in pseudomonas aeruginosa pa01. 2021;67(8).

23. Rabbi MF, Akter SA, Hasan MJ, Amin AJB, insights b. In silico characterization of a hypothetical protein from Shigella dysenteriae ATCC 12039 reveals a pathogenesis-related protein of the type-VI secretion system. 2021;15:11779322211011140.

24. Yu NY, Wagner JR, Laird MR, Melli G, Rey S, Lo R, et al. PSORTb 3.0: improved protein subcellular localization prediction with refined localization subcategories and predictive capabilities for all prokaryotes. 2010;26(13):1608–15.

25. Sivakumar K, Balaji S, Sciences GJJoC. In silico characterization of antifreeze proteins using computational tools and servers. 2007;119:571–9.

26. Marchler-Bauer A, Derbyshire MK, Gonzales NR, Lu S, Chitsaz F, Geer LY, et al. CDD: NCBI’s conserved domain database. 2015;43(D1):D222–D6.

27. El-Gebali S, Mistry J, Bateman A, Eddy SR, Luciani A, Potter SC, et al. The Pfam protein families database in 2019. 2019;47(D1):D427–D32.

28. Jones P, Binns D, Chang H-Y, Fraser M, Li W, McAnulla C, et al. InterProScan 5: genome-scale protein function classification. 2014;30(9):1236–40.

29. Benkert P, Tosatto SC, Schomburg DJPS, Function,, Bioinformatics. QMEAN: A comprehensive scoring function for model quality assessment. 2008;71(1):261–77.

30. Wiederstein M, Sippl MJJNar. ProSA-web: interactive web service for the recognition of errors in three-dimensional structures of proteins. 2007;35(suppl_2):W407-W10.

31. Tian W, Chen C, Lei X, Zhao J, Liang JJNar. CASTp 3.0: computed atlas of surface topography of proteins. 2018;46(W1):W363–W7.

32. DeLano WLJCNPC. Pymol: An open-source molecular graphics tool. 2002;40(1):82–92.

